# Phosphoproteomic analysis of chondrocytes after short-term exposure to inorganic polyphosphate

**DOI:** 10.1101/2021.07.30.454470

**Authors:** Uros Kuzmanov, Rahul Gawri, Alena Zelinka, Keith A Russell, Shin-Haw Lee, Anthony Gramolini, Rita Kandel

## Abstract

Osteoarthritis is a debilitating disease of the joint that affects over 230 million people worldwide. Currently there are no treatments that slow the progression of this disease. For these reasons, new biological treatment options are currently being explored. Inorganic polyphosphates are naturally occurring biological molecules that have an anabolic effect on chondrocytes grown *in vitro* in the presence of Ca^2+^. We hypothesized that when examining significant changes in protein phosphorylation, key candidates would emerge that could help to elucidate the anabolic effects of polyphosphate on chondrocytes. Therefore, we conducted a large-scale quantitative proteomic and phosphoproteomic study of bovine primary articular chondrocytes after 30-minute treatment with inorganic polyphosphate and Ca^2+^. Mass spectrometry identified more than 6000 phosphorylation sites on ∼1600 chondrocyte phosphoproteins while proteomic analysis detected approximately 4100 proteins. Analysis of the data revealed a swift and dynamic response to polyphosphate after 30 minutes. What emerged from the list of proteins most affected by the treatment were proteins with key roles in chondrogenesis including TNC, IGFBP-5, and CTGF, indicating that polyphosphate plays an important role in chondrocyte metabolism. This phosphoproteome serves as a meaningful resource to help elucidate the molecular events that contribute to extracellular matrix production in cartilage.

## Introduction

Osteoarthritis (OA), the most common form of arthritis, is a progressive disease of the joint with no disease-modifying treatments currently available. Early disease can be treated with anti-inflammatory drugs, exercise, weight loss, and/or physiotherapy, but as it becomes more advanced, surgical intervention, such as joint replacement, may be necessary. Some patients require revision surgeries, a procedure that is more common in younger patients.^1^ Thus, there is an unmet need to develop biological therapeutic approaches to the treatment of this disease to arrest progression. Alternative therapeutic approaches currently under investigation include cell or tissue implant based therapies, ^2^ and developing naturally occurring biological molecules that induce anabolic cell responses in articular chondrocytes. ^3^ An example of one such agent is inorganic polyphosphate (polyP). ^4^

Inorganic polyphosphates (polyP), found in every form of life, are linear biopolymers made up of few to hundreds of orthophosphate (PO_4_) subunits connected by high-energy phosphoanhydride bonds. Study of this macromolecule, primarily in microorganisms, has revealed a diverse variety of functions ranging from serving as an energy source akin to ATP to chelating metal cations. ^5^ Over the last 25 years, interest in polyP function in mammalian cells has increased as more sensitive methods to analyze it have been developed and as we appreciate its role in critical metabolic and signaling processes. PolyP is no longer thought to be a “fossil molecule”. Functionally, in mammals, it has been associated with bone formation, ^6^ hemostasis, ^7^ astroglial signaling, ^8^ and energy metabolism. ^9^ Interestingly, the chain length and concentration of polyP range widely across tissues, cells, and even subcellular locations. ^10^ Our group has shown that the most effective polyphosphate chain length and dose required to stimulate anabolic processes *in vitro* is cell type-dependent. PolyP chain length of 45 phosphate residues (polyP-45) at a concentration of 1 mM was optimal in stimulating matrix accumulation by chondrocytes grown in 3D cultures. ^4^ In contrast, a chain length of 22 phosphate residues (polyP-22) at 0.5 mM had the greatest anabolic effect on nucleus pulposus cells. ^11^ Until recently, the mechanism by which polyP induces anabolic responses in chondrocytes was unknown. Recently our group demonstrated that polyP regulates calcium influx in chondrocytes through modulation of voltage-gated and mechanosensitive calcium ion channels, which stimulated glycosaminoglycan accumulation. ^12^ Further, following treatment with polyP and Ca^2+^, polyP was detected in intracellular organelles including the nucleus within 30 minutes. This suggests that polyP could have multiple functions within the chondrocyte, such as increasing intracellular calcium levels.

The reversible covalent attachment of phosphate groups is one of the most common post-translational modifications found on proteins, and it is widely recognized as a key regulatory and signaling mechanism in a variety of cellular processes. Protein kinases are the enzymes that catalyze the transfer of a phosphate group from ATP to serine, threonine, and tyrosine residues on target proteins. ^13^ There are approximately 500 putative protein kinases in the human genome, ^13^ and anecdotal evidence suggests that a third of all proteins are phosphorylated. Mass spectrometry (MS) techniques allow for global identification and even quantification of phosphorylation sites. Systems level bioinformatic filtering, identification, and analysis of signaling pathways ultimately allow for the identification of altered cellular mechanisms and signaling pathways between different experimental conditions.

In order to attain an unbiased global view of the effect of polyP on bovine chondrocytes, we conducted a large-scale quantitative proteomic and phosphoproteomic study of chondrocytes after short-term treatment with polyP and Ca^2+^. Calcium was necessary for the anabolic effect of polyP. We hypothesize that when examining significant changes in protein phosphorylation, key candidates would emerge that could help to elucidate the anabolic effects of polyP on chondrocytes. MS identified in excess of 6000 phosphorylation sites on approximately 1600 bovine chondrocyte phosphoproteins in addition to a further ∼4100 bovine proteins identified by proteomic analysis. Owing to the utilization of TMT-based isobaric labelling of (phospho)peptides, a majority of identified phosphorylation sites/proteins were reliably quantified. Bioinformatic analysis revealed dozens of altered cellular processes and signaling pathways caused by polyP treatment of bovine chondrocytes in culture suggesting an important role for polyP in affecting cell metabolism and biology.

## Experimental procedures

### Cell isolation

Metacarpophalangeal joints were obtained from 9-12 month old calves from the local abattoir. Knee joint capsules were opened under aseptic conditions, and articular cartilage tissue was harvested. Cartilage from 3 legs was pooled. The tissue was minced, sequentially digested with 0.25% protease (Sigma, St. Louis, MO, USA) in Ham’s F12 media (Wisent, St. Bruno, QC, Canada) supplemented with 1% antibiotic-antimycotic solution (Wisent, Saint-Jean-Baptiste, QC, Canada) for 1 hour, washed twice with sterile phosphate buffered saline (PBS), and digested overnight with 0.1% Collagenase A (Roche Diagnostics, Mannheim, Germany) in Ham’s F12 media supplemented with 1% antibiotic-antimycotic solution at 37°C in 5% CO_2_. ^14^ The following day, the digest was passed through a 40 µm cell strainer and washed three times with Ham’s F12 medium, and the total cell number was determined by hemocytometer using Trypan Blue Solution (Gibco, Burlington, ON, Canada).

### Cell seeding and treatment with polyP

Chondrocytes were seeded in T-175 cell culture flasks (Corning®, Sigma-Aldrich, Oakville, ON, Canada) at a density of 50,000 cells/cm^2^ (∼9 million cells per flask) in Ham’s F12 medium supplemented with 5% fetal bovine serum (FBS, Wisent, St. Bruno, QC, Canada) and 1% antibiotic-antimycotic solution at 37°C in 5% CO_2_. The following day, all flasks were washed with serum-free Ham’s F12 medium three times for 5 minutes each and then serum-starved in Ham’s F12 medium for 1 hour before the experiment was initiated.

### Experimental design and statistical rationale

The cells were then either co-treated with 1.5 mM Ca^2+^ (diluted from filter-sterilized 1M CaCl_2_ solution; Sigma-Aldrich, Oakville, ON, Canada) and 1 mM polyP-45 (average chain length 45 P_i_ units; Sigma-Aldrich, Oakville, ON, Canada) in Ham’s F12 medium for 30 minutes or medium alone (control). The control cells were either collected immediately after the media change or 30 minutes later. Cells collected right after feeding were to control for the effect of the media change. Two technical replicates were performed at the MS level (i.e. samples were run twice on the mass spectrometer). No randomizations were used except for the volcano plot where default settings were used (T-test, 250 randomizations) as previously described. ^15^ The use of the TMT isobaric tags limited us to 10 total channels (i.e. samples) available for increased quantitative reliability through the cumulative decrease in technical error in sample processing and MS experimental steps. This led us to use the t-test and ANOVA for performing pairwise and multiple sample group comparisons, respectively.

### Cell harvesting

At the end of the incubation period, cell culture media was removed and cells were harvested using a cell scraper with 800 μl cell lysis buffer containing 8M urea solution (reagent grade, 98%, Sigma-Aldrich, Oakville, ON, Canada), supplemented with protease inhibitor (complete™, Mini, EDTA-free Protease Inhibitor Cocktail, 1 tablet per 10 mL lysis buffer, Sigma-Aldrich, Oakville, ON, Canada) and phosphatase inhibitors (1 mM activated sodium orthovanadate and 1 mM sodium fluoride, Sigma-Aldrich, Oakville, ON, Canada) on ice. The cell extracts were collected and flash-frozen in an alcohol bath (100% ethanol on dry ice). ^16^ Protein concentration of the extracts was estimated using a BCA protein assay kit (Pierce™ BCA Protein Assay Kit, ThermoFisher Scientific, Oakville, ON, Canada) as per supplier’s instructions. The cell extract solution was diluted by a factor of four to a final urea concentration of 2M so the BCA protein assay kits could be used.

### Sample preparation for mass spectrometry

A total of 100μg of protein from each sample was subjected to trypsin digestion as previously described. ^16^ Briefly, each sample was reduced by 2.5mM DTT for 60 minutes at room temperature and alkylated by 5mM iodoacetamide for 30 minutes at room temperature in the dark. Samples were diluted 10X in 50 mM ammonium bicarbonate after which 4 µg of sequencing-grade trypsin (Promega, Madison, WI, USA) was added before overnight incubation at 37°C. The reaction was stopped by the addition of formic acid (to 1% in solution). Resulting tryptic peptides were desalted using C18 TopTip (Glygen, Petersborough, ON, Canada) as per manufacturer’s instructions, dried to completion by SpeedVac (ThermoFisher Scientific, Oakville, ON, Canada), labelled with TMT10-plex reagents as described by manufacturer (ThermoFisher Scientific, Oakville, ON, Canada), and combined following the labelling procedure. The combined sample was again dried to completion by SpeedVac and resuspended in 80% acetonitrile (ACN) with 0.1% trifluoroacetic acid (TFA).

### HILIC fractionation and phosphopeptide enrichment

HILIC chromatography was performed on an Agilent 1200 HPLC system connected to a 2.0 × 150 mm 5-μm particle TSKgel Amide-80 column (Tosoh Biosciences, San Francisco, CA, USA) as previously described. ^16^ The TMT-labelled sample was loaded onto the column at a flow rate of 250 μL/min with 80% buffer A (98% ACN, 0.1% TFA) and 20% buffer B (2% ACN, 0.1% TFA). The liquid chromatography gradient was as follows: 3 min loading with 20% buffer B, a linear gradient from 20–40% buffer B between 3 and 30 min, a 3 min gradient of 40–100% buffer B, continuous flow of 100% buffer B for 5 min, a gradient of 100–20% buffer B for 2 min, and finally 20% buffer B for 10 min. Eluted fractions were collected in 1.5 ml tubes at 2 min intervals and later combined (first four fractions were analyzed individually and the remaining 21 collected fractions were combined sequentially in groups of 3) to form 11 distinct fractions which were lyophilized to dryness by SpeedVac. 10% of each fraction was kept for global proteomic analysis. Phosphopeptide enrichment was performed employing TiO_2_-coated Mag Sepharose beads (GE Healthcare, Mississauga, ON, Canada) as per manufacturer’s protocol. Briefly, the dried peptide mixture from each HILIC fraction was resuspended in 200 μl of binding buffer (1M glycolic acid, 80% ACN, 5% TFA), incubated with TiO_2_ beads for 30 min at room temperature before the supernatant was removed. The beads were then washed once with binding buffer and three times with wash buffer (80% ACN, 1% TFA); phosphopeptides were eluted in 5% ammonium hydroxide and dried to completion by SpeedVac prior to LC-MS analysis.

### Liquid chromatography-mass spectrometry analysis

Phosphopeptide mixtures from each HILIC fraction were individually loaded onto a reverse phase Thermo Acclaim PepMap pre-column (2 cm length, 75 μm diameter, 3 μm C18 beads) and separated on a Thermo PepMap RSLC C18 analytical column (50 cm length, 75 μm diameter, 2 μm C18 beads) connected to an Easy-nLC 1200 system (Thermo). The nanoflow gradient was composed of buffer A (5% ACN, 0.1% formic acid) and buffer B (85% ACN, 0.1% formic acid). Equilibration was performed with 100% buffer A on both the pre- (20 μl) and analytical columns (3 μl) prior to each injection. The 3 hour nanoflow gradient (220 nl/min flow rate) was as follows: 5-35% buffer B for 156 min, 35-100% buffer B for 9 min, 100% buffer B for 15 min.

Peptides and phosphopeptides were directly ionized by an EasySpray ion source (ThermoFisher Scientific)) into an Q Exactive HF mass spectrometer (ThermoFisher Scientific)). 15 MS2 data-dependent scans were acquired per single MS1 full scan mass spectrum using profile mode with HCD fragmentation. Full scans were performed at the 120,000 resolution setting, maximum injection time of 50 ms, ion packet setting of 3 × 10^6^ for automatic gain control, and a 350 to 1450m/z range. MS2 scans were performed at 60,000 resolution, ion packet setting of 1 × 10^5^ for automatic gain control, maximum injection time of 100ms, fixed first mass at 100m/z, 1.2 m/z isolation window and using 32% normalized collision energy. The dynamic exclusion range was set to 20 s and unassigned, 1+, and parent ions with charge states higher than 6 were excluded from MS2 analysis. Liquid chromatography and MS settings and methods were identical for proteomic (unenriched) and phosphoproteomic (phosphopeptide enriched) samples/fractions.

Resulting RAW files were searched using MaxQuant (version 1.5.5.1; http://maxquant.org/) with “Reporter ion MS2” 10plex TMT settings using an ENSEMBL Bos taurus protein database allowing for three missed trypsin cleavage sites and a reporter ion tolerance of 0.01. Variable modifications were set for phosphorylation at serine, threonine, and tyrosine residues, N-terminal acetylation, and methionine oxidation. Carbamidomethylation of cysteine residues was set as a fixed modification. First and second search ion tolerances were set at 20 and 4.5 ppm, respectively. Candidate (phospho)peptide, protein, and phosphorylation site identifications were filtered based on a 1% false discovery rate threshold based on searching of the reverse sequence database. The MS proteomics data have been deposited to the ProteomeXchange Consortium (http://proteomecentral.proteomexchange.org) via the PRIDE partner repository with the data set identifier PXD014204.

### Data analysis

MaxQuant search data was analyzed by Perseus (version 1.6.0.7) and Bioconductor packages in R. ^15, 17^ All proteomic and phosphoproteomic data were log_2_-transformed and quartile normalized by “width adjustment” in Perseus. Identified phosphorylation sites were filtered based on a 0.7 site identification probability, and quantitative values for each site were combined and collapsed only for phosphopeptides carrying the same number of modifications. Phosphoproteomic and proteomic data sets were merged at the gene/protein level. Duplicate entries were removed. To probe altered pathways, the merged data set was analyzed by statistical overrepresentation test or gene set enrichment analysis (GSEA). PANTHER statistical overrepresentation test ^18^ was performed on significantly up or downregulated genes identified by multi-way ANOVA. For GSEA, gene sets were obtained from a custom human “all pathways” database (http://download.baderlab.org/EM_Genesets/) carrying annotated gene ontology biological process terms and curated pathways ^19^ and were required to have between 10 to 300 associated components. False discovery rate (FDR, q-value) was calculated based on 1000 gene set permutations. The “classic” GSEA enrichment statistic was utilized and t-test significance levels between pairs of experimental conditions were used to rank gene/protein IDs.

### Kinase motif analysis

Identified phosphorylation sites (with surrounding sequences) that were significantly altered between the experimental conditions (p < 0.05) were searched for consensus motifs using the Motif-X algorithm. ^20^ Central residues were set as S or T with the “MS/MS” foreground format searched against the human proteome. Window width was set to 13 and significance was set to 0.000001. Identified consensus sequences were then searched for kinases known to target these motifs using PhosphoMotif Finder. ^21^

### STRING analysis

Protein-protein associations for phosphoproteins in which there were significant changes in phosphorylation between experimental conditions (p < 0.05) were identified using the STRING database (version 10.5). ^22^

## Results

### Phosphoproteomic analysis

Global quantitative mass spectrometry (MS)-based proteomic and phosphoproteomic profiling was performed to map protein and phosphorylation site changes resulting from polyP+Ca^2+^ treatment of isolated bovine chondrocytes in culture. The samples were processed with attention to maintaining phosphorylation site integrity through use of phosphatase inhibitors, low temperature, and protein denaturing conditions ensuring minimal phosphatase activity. A TMT10-plex isobaric (phospho)peptide labelling strategy was applied to minimize technical error associated with multiple experimental steps required in phosphoproteomic analyses and provide robust quantification. HPLC hydrophilic interaction liquid chromatography (HILIC) was utilized to pre-fractionate peptides and phosphopeptides resulting from tryptic digestion of analyzed protein mixtures to ensure in-depth coverage. Additionally, titanium dioxide (TiO_2_) affinity enrichment chromatography was performed to ensure confident identification of phosphopeptides, due to their low abundance and susceptibility to ion suppression in MS analyses. Quantification was performed using extracted TMT ion reporter intensities recorded in high resolution MS2 scans on the precision Thermo Q Exactive HF orbitrap instrument. Total protein levels were measured in parallel (Figure 1A).

**Figure 1.**
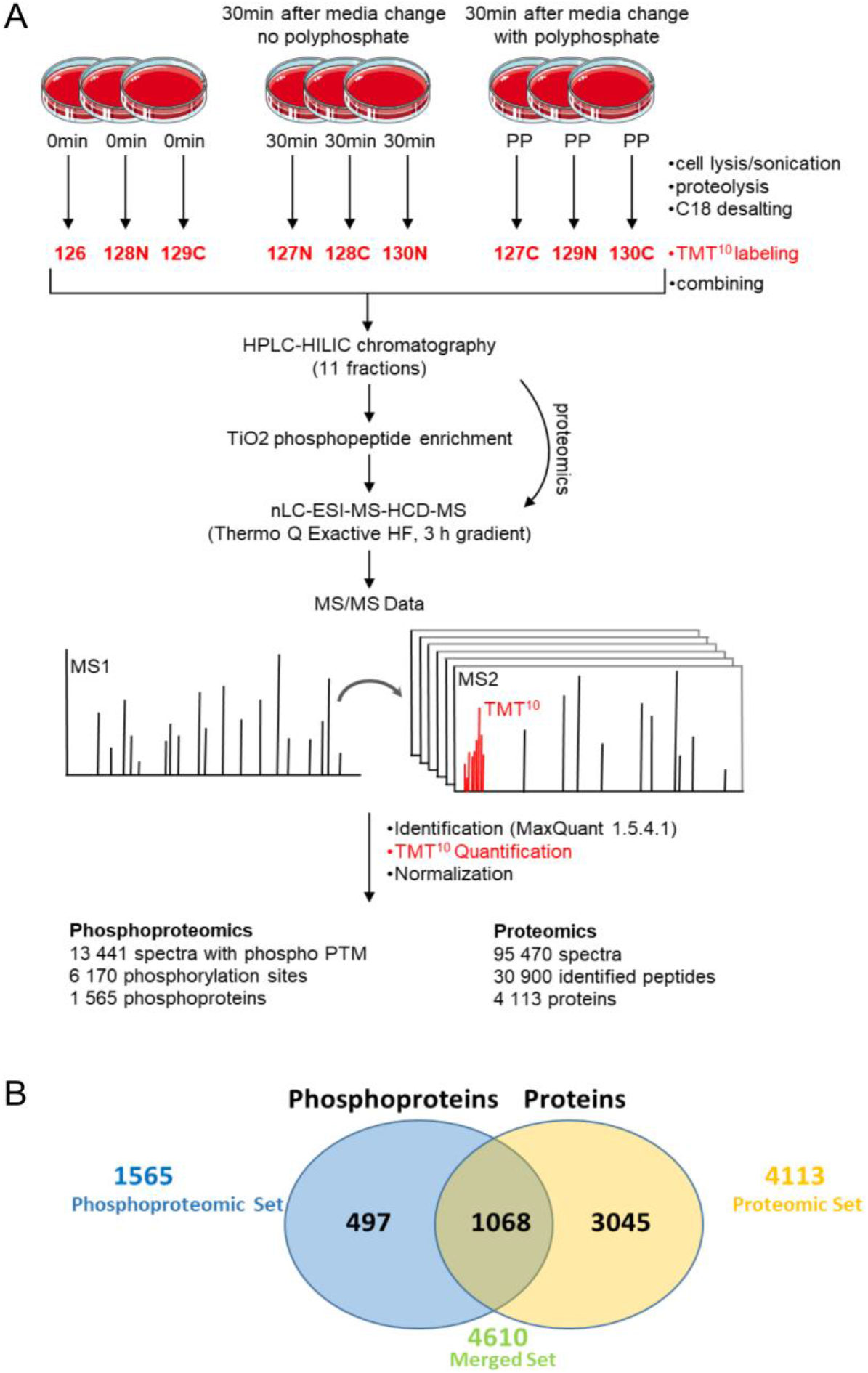
(A) Overview of experimental procedures and results of the TMT-based quantitative (phospho)proteomic analysis of Ca+ polyphosphate treated bovine chondrocytes and relevant controls accounting for media replacement. (B) Protein ID level coverage and overlap between phosphoproteomic and proteomic analyses.

### Data summary

As a result of the phosphoproteomic analysis totaling 20 mass spectrometry runs (10 HILIC fractions performed in technical duplicates), 13441 MS2 spectra were assigned to phosphopeptide sequences (at 1% FDR). These peptides mapped to 1565 chondrocyte phosphoproteins identifying 5050 distinct phosphorylation sites (∼82% of which could be classified as high confidence with a 0.7 site localization probability, Figure 1B). The vast majority of identified phosphorylation sites were located on 3697 serine residues, with 408 and 26 mapping to threonine and tyrosine, respectively. Additionally, 4113 proteins were identified in parallel utilizing the proteomics component of the workflow allowing, in total, the combined identification of 4610 bovine chondrocyte proteins. The majority of identified proteins and phosphorylation sites (>95%) had associated quantitative values across all samples owing to the use of TMT10-plex isobaric tags for the labelling of (phospho)peptides. This is not the case with label-free approaches where the proportions of missing values between experimental samples for individual phosphopeptides/proteins can account for a large proportion of assigned identifications. Both proteomic and phosphoproteomic data sets showed a normal distribution of values and high levels of correlation between samples (Supplementary Figures 1 and 2).

### Quantitative comparison

Using TMT reporter ion measurements, 451 phosphorylation sites on 265 proteins were found to be significantly altered between the polyphosphate treated samples and 30 minute controls (Figure 2A) based on a two-tailed Student’s t-test (p < 0.05), permutation based FDR (q < 0.05), and artificial within group variance (S0 > 0.1). A majority of these phosphorylation sites (59.6%) were downregulated as result of polyphosphate treatment of cells. Comparatively, the expression of 255 proteins was shown to be significantly altered as a result of polyP+Ca^2+^ co-treatment (Figure 2A, q<0.05). Among these, proteins were noted that have been identified to play a role in chondrocyte biology including CTGF (connective tissue growth factor) and TNC (Tenascin C) (Figure 2B).

**Figure 2.**
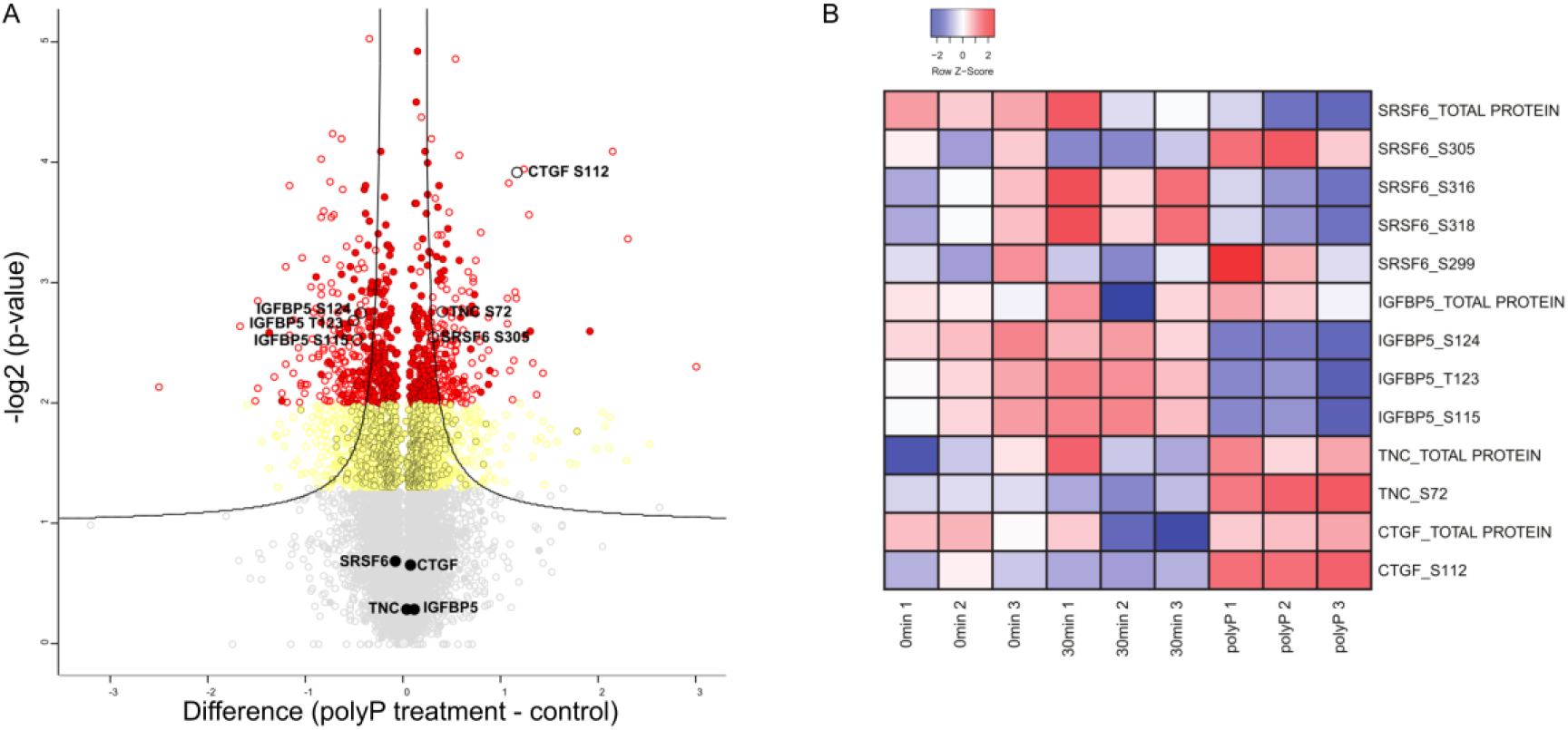
(A) Volcano plot showing the differences between polyP-treated and 30 minute control (media change alone) merged data sets (all protein and phosphorylation site quantitative information). IDs are represented by filled (proteins) and open (phosphorylation site) circles. The x-axis represents the difference in means. Student’s t-test significantly altered proteins/phosphorylation sites are highlighted in yellow (p<0.05) and red (p<0.01). The highlighted lines represent an FDR cutoff (q<0.05) at an S0 = 0.1 artificial within group/sample type variance. Selected significantly altered phosphorylation sites showing no change at the protein level are highlighted. (B) Z-scored value distributions of phosphorylation sites and related protein levels of candidates relevant to chondrocyte biology.

### Unsupervised global analysis

For further statistical analyses, the proteomic and phosphoproteomic data (Supplementary Data S1) were independently normalized (see Methods) and then combined into a single merged set with multiple gene/protein level entries for phosphoproteins (i.e. individual entries for protein level expression and each phospho site identified). Sites that were identified on singly or multiply phosphorylated phosphopeptides were also treated as separate entries in the merged data set. Unsupervised hierarchical clustering of the merged data set demonstrated clear segregation between the 0 and 30 minute (control for media change) experimental time points in addition to their distinction for the polyphosphate treated samples (Figure 3A). Similarly, analyzing the global merged data utilizing principal component analysis (PCA), a similar trend of clear differentiation between the three experimental groups was observed (Figure 3B).

**Figure 3.**
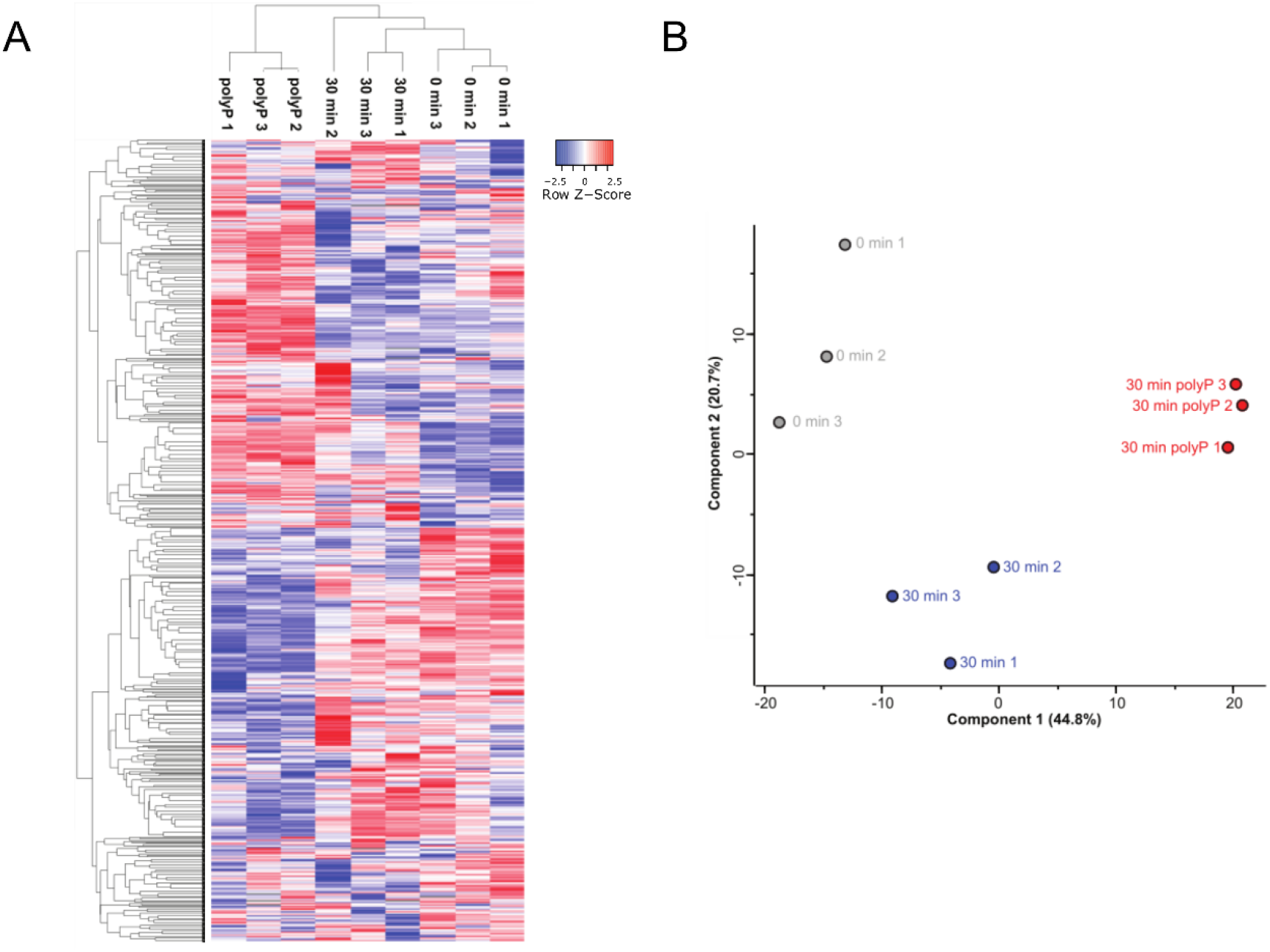
(A) Unsupervised hierarchical clustering heat map of all protein and phosphorylation site z-score normalized values across all three experimental conditions. (B) Principal component analysis of all protein and phosphorylation site normalized values across all three experimental conditions.

### Kinase motif analysis

Motif-X analysis identified 11 motifs (listed in Table 1, Supplementary Data S1) targeted within 30 minutes of polyP+Ca^2+^ treatment. Kinases involved in phosphorylation of these motifs include ERK1/2, CAMK2, GSK-3, PKA, PKC, GPCRK1, CK2, CK1, and CDK5.

**Table 1.** Consensus motifs based on centered 13 residue sequences surrounding phosphorylation sites found to be significantly altered between the polyphosphate treated cells and 30 minute media change control. Motif-X score and fold increase over the normal occurrence of consensus sequences in the proteome is also provided.

| Motif | Motif Score | Fold Change | Targeting Kinase |
| --- | --- | --- | --- |
| ...R..SP.P... | 36.77 | 20.71 | PKA, PKC, CAMK2, GSK-3, ERK1, ERK2, CDK5 |
| ..SP..SP..... | 38.05 | 16.11 | GSK-3, ERK1, ERK2, CDK5 |
| .....SD.E... | 28.98 | 11.13 | CK2 |
| ...RS.S..... | 32 | 6.42 | PKA, PKC, CAMK2 |
| .....SSP.... | 32 | 5.63 | GSK-3, ERK1, ERK2, CDK5 |
| .....S...SP. | 31 | 5.61 | GPCRK1 |
| .....SEE.... | 22.77 | 5.04 | GPCRK1, CK1 |
| .....S..SP.. | 26.66 | 4.96 | GPCRK1, CK1 |
| .....TP..... | 16 | 4.71 | GSK-3, ERK1, ERK2, CDK5 |
| .....SP..... | 16 | 2.83 | GSK-3, ERK1, ERK2, CDK5 |
| ...R..S..... | 16 | 2.35 | PKA, PKC, CAMK2 |

### STRING analysis

STRING analysis identified potential interactions amongst phosphoproteins involved in RNA transport, mRNA surveillance, spliceosome, and endocytosis processes. These pathways are listed in Table 2 (Supplementary Data S2).

**Table 2.** Phosphoproteomic pathway analysis of 408 unique proteins based on significant changes in abundance of 1175 phosphosites in polyphosphate-treated versus non-treated bovine chondrocytes identified by the online STRING database.

| Pathway | Protein count | False discovery rate | Matching proteins |
| --- | --- | --- | --- |
| RNA transport | 14 | 0.000713 | ACIN1, EIF4B, EIF4EBP1, EIF4G3, EIF5B, FMR1, NUP153, NUP214, PNN, RANBP2, RNPS1, SEC13, SRRM1, UPF1 |
| mRNA surveillance pathway | 9 | 0.00461 | ACIN1, CPSF6, CSTF3, PCF11, PNN, PPP2R5E, RNPS1, SRRM1, UPF1 |
| Spliceosome | 11 | 0.00461 | ACIN1, DDX42, DDX46, PRPF38B, SF3B1, SF3B2, SFRS13A, SRSF4, SRSF6, SRSF9, U2AF2 |
| Endocytosis | 13 | 0.00897 | AGAP3, DNM1, FAM125A, GIT2, IQSEC1, LDLRAP1, NEDD4, PARD3, PDGFRA, PML, RABEP1, SH3KBP1, VPS4B |

### Pathway analysis

A protein/gene ID level statistical overrepresentation test was performed ^18^ on significantly (multi-way ANOVA, p<0.05) up- and downregulated proteins and phosphorylation sites resulting from polyP treatment (Figure 4A, Supplementary Data S3). The majority of annotations associated with signaling pathways were identified based on (phospho)proteins/phosphorylation sites found to be downregulated as a result of polyP treatment. Conversely, several annotations associated with kinase inhibition (highlighted in Figure 4A) were found to be overrepresented in the group of upregulated data suggesting a general inhibition of kinase activity and suppression of signaling pathways.

**Figure 4.**
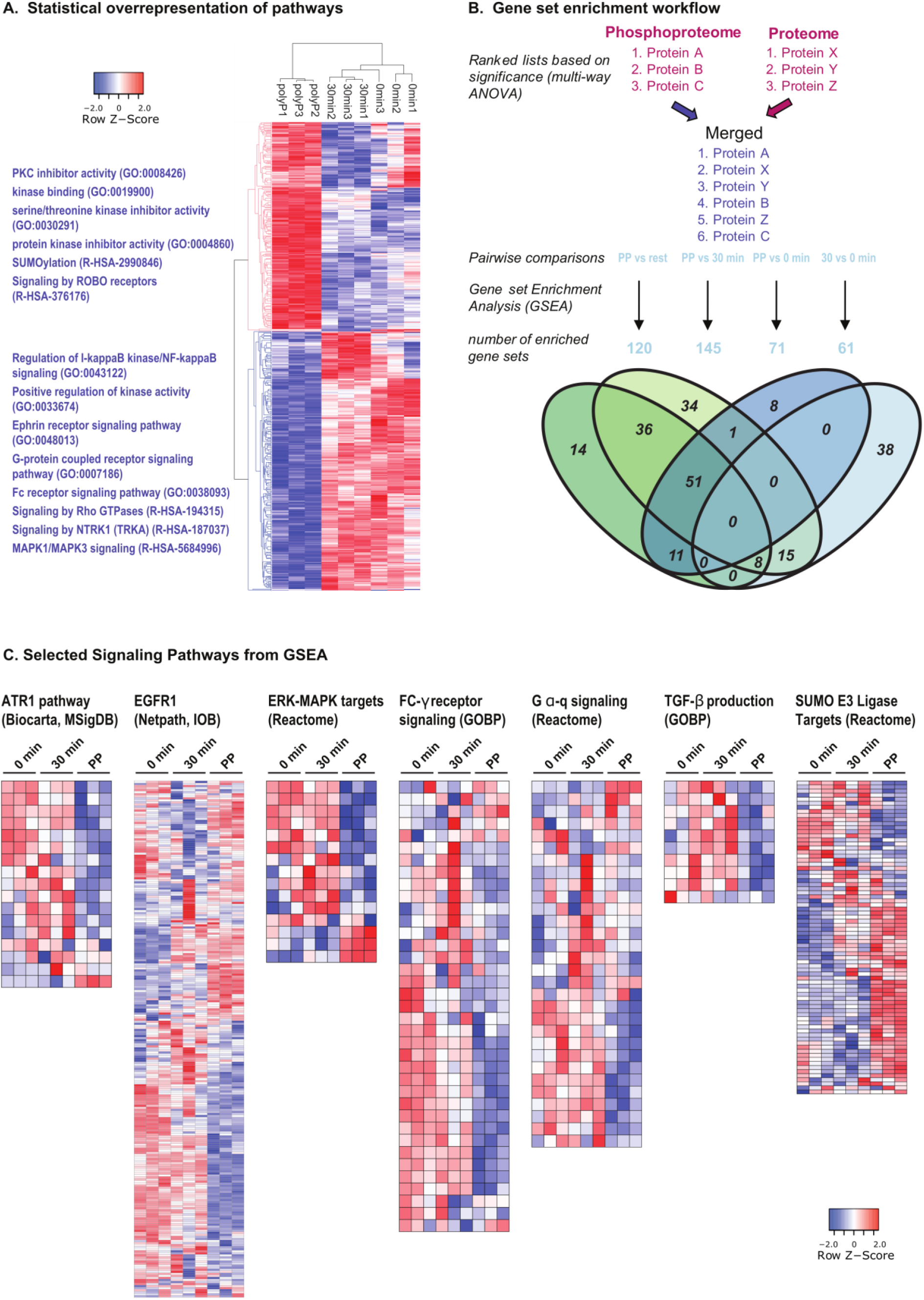
(A) Hierarchical clustering heat map of multi-way ANOVA (p<0.05) significant proteins and phosphorylation sites (z-score normalized values). Selected annotations associated with up- and downregulated proteins (statistical overrepresentation test) are highlighted on the left side. (B) Overview of the workflow for gene set enrichment analysis. Venn diagram shows overlap of significantly enriched gene sets/annotations (q<0.05) in pairwise comparisons of control experimental conditions and polyP treated chondrocytes. (C) Heat maps show the most significantly altered gene/protein IDs (indicating significant changes in phosphorylation sites or protein level) from selected signaling pathway gene sets.

To further leverage the quantitative information associated with identified proteins and phosphorylation sites, gene set enrichment analysis (GSEA) was performed in pairwise fashion comparing all experimental conditions to polyP treated cells (Figure 4B, Supplementary Data S4, Supplementary Figures 4A and 4B). In the case where gene/protein IDs were listed multiple times, only the most significant entry for each gene/protein ID was used. Similar to the overrepresentation test, signaling pathways associated with dynamic protein phosphorylation states were identified as altered in comparisons of polyP+Ca^2+^ co-treated to non-treated conditions (media change and no media change) individually or combined. For example, MAPK/ERK/ATR1 signaling, EGFR pathways, and G-α-q signaling were found to be downregulated as a result of exposure to polyP (Figure 4C). Conversely, a gene set containing proteins associated with targets of SUMO E3 ligases was found to be upregulated based on the expression levels of member proteins.

## Discussion

Identification of pivotal molecular events contributing to synthesis of cartilage macromolecules, their degradation, and homeostasis is necessary if effective treatments are to be developed for osteoarthritis. Polyphosphates are bioactive molecules that have anabolic effects on cartilage matrix synthesis and accumulation. ^4^ In this study, we report a global phosphoproteomic analysis of the effect of short-term polyP+Ca^2+^ exposure on chondrocytes. We quantified 5050 phosphorylated peptides on 1565 proteins, of which there were significant changes in abundance in 1175 phosphosites mapping to 408 unique proteins.

Analysis of these phosphoproteins using the online STRING database revealed networks of proteins involved in RNA and protein processing including RNA transport, mRNA surveillance, spliceosome, and endocytosis pathways (Table 2). To see such rapid activation of transcriptional and translational machinery after only 30 minutes of polyP+Ca^2+^ treatment is indicative of a quick and potent general effect. To support this, included in the top 75 differentially phosphorylated proteins are eukaryotic translation initiation factor 4B (EIF4B) and, a key kinase of EIF4B, ribosomal protein S6 kinase alpha-1 (RPS6KA1). RPS6KA1 is in turn activated by ERK1/2, ^23^ both of which appear repeatedly in the kinase motif analysis (Table 1). Time-course experiments tracking EIF4B and RPS6KA1 performed by Shahbazian et al. showed rapid phosphorylation kinetics within the 30 minute time frame reported here. ^23^ Phosphorylated EIF4B (on serine 422) has been shown to activate helicase EIF4A and complex with EIF3, which in turn promotes protein synthesis. ^23, 24^

We identified some candidates within the top 75 unique proteins that may contribute to the stimulation of matrix production based on previous literature. One example is tenascin-C (TNC), which is active in early chondrogenesis ^25^ and chondroprotective in a murine OA model. ^26^ To substantiate the presence of TNC, serine/arginine-rich splicing factor 6 (SRSF6), a protein associated with the alternative splicing of TNC, ^27^ also appears in this list of differentially phosphorylated proteins. Phosphorylation of TNC has not been thoroughly investigated. However, the same phosphorylated serine (S72) that we identified is the most abundantly referenced in high throughput studies (searching TNC at phosphositeplus.org). One high throughput study in particular identified the kinase Fam20C as responsible for the phosphorylation of TNC, in addition to the phosphorylation of the majority of secreted proteins. ^28^

Another phosphoprotein identified, insulin growth factor binding protein 5 (IGFBP-5), is also phosphorylated by Fam20C. ^28^ IGFBP-5 is a matricellular protein which enhances both proliferation and differentiation of chondrocytes through regulation of IGF. ^29, 30^ In our data, we observed IGFBP-5 in polyP+Ca^2+^-treated chondrocytes to be less phosphorylated than in the controls. However, it is unclear how this change would affect IGF bioactivity or the IGF-independent actions of IGFBP-5. Interestingly, Graham et al. have shown that phosphorylation of IGFBP-5 decreases binding affinity to the glycosaminoglycan heparin sulfate, but not IGF-I or IGF-II. ^31^ Thus, less phosphorylation in the case of polyP exposure may result in stricter binding of IGFBP-5 to other glycosaminoglycans in the extracellular matrix and provide a store of IGF for cartilage synthesis.

Connective tissue growth factor (CTGF) is another matricellular protein with binding affinity to IGF as it contains an IGFBP domain. CTGF is essential in development as CTGF-deficient mice are born severely chondrodysplasic. ^32^ CTGF is required for cartilage extracellular matrix production, organization, and remodeling. ^33, 34^ It is also secreted by nucleus pulposus cells, and *in vitro* studies suggest that it enhances proteoglycan gene expression. ^35^ One study demonstrated that overexpression of CTGF under control of the type II collagen promoter in aging transgenic mice had a chondroprotective effect. ^36^ Proteins associated with chondrogenesis, SOX9 and TGF-β, have been indicated as direct regulators of CTGF expression. ^37-39^ However, regulation of CTGF through phosphorylation has not been extensively studied, and there are no previous reports of the phosphosite serine 112 seen in our data even in high throughput studies. ^40^

The data suggests that these phosphoproteins alone or in concert may be responsible for the anabolic effect of polyP+Ca^2+^ co-treatment. While phosphorylation of a residue may not translate into altered physiological function, it has been suggested that conservation of the position and flanking residues of a phosphorylated site across species might be indicative of its relevance. ^41^ Of the phosphorylation sites discussed here, all meet these criteria for relevance with the exception of CTGF. Nevertheless, owing to the significant relation of CTGF to chondrogenesis, it is likely that it also warrants further study.

It was not unexpected to see so many phosphorylated proteins after 30 minute treatment as others have shown that most phosphosites are phosphorylated after 10 minutes of stimulation. ^42^ Indeed, we showed previously that polyP is able to enter into chondrocytes and organelles within 30 minutes in the presence of calcium. ^12^ One of the limitations of mass spectrometry is that polyphosphorylation of proteins cannot be detected due to the variable chain lengths of polyP (i.e. search algorithms used in MS-based proteomic analyses require a clear specified mass change on identified phosphopeptides) and its phosphoramidate bond with the lysine residue, which is too unstable for the conditions of the method. ^43^

In conclusion, we provide an unbiased comprehensive phosphoproteome profile of the effect of polyP on chondrocytes. Analysis of the data exposed a swift and dynamic response to the treatment after only 30 minutes. What emerged from the proteins most affected by the treatment were key proteins known for their role in chondrogenesis including TNC, IGFBP-5, and CTGF. Going forward, this phosphoproteome will be an important resource that may help untangle the molecular events that lead to cartilage anabolism when treated with polyP.

## Supporting information

all proteomic and phosphoproteomic data

STRING pathway analysis

Data and Results for Statistical overrepresentation in significant hits

GSEA results - significantly enriched gene sets

## Acknowledgements

This project was funded by the Translational Biology and Engineering Program Seed Fund to A.O.G. and CIHR MOP 126111 to RK. U.K. received a Ted Rogers Centre for Heart Research Fellowship. S-H.L was supported by a NSERC Postgraduate Scholarship, an Ontario Graduate Scholarship, and a Ted Rogers Centre for Heart Research Doctoral Fellowship. RG was supported by an NSERC Post-doctoral Fellowship.

## Data availability

The MS proteomics data have been deposited to the ProteomeXchange Consortium (http://proteomecentral.proteomexchange.org) via the PRIDE partner repository with the data set identifier PXD014204 (for purposes of the review: username: and password: gr1Y5VLe.).

## Abbreviations

ACN: acetonitrile
CTGF: connective tissue growth factor
EIF4B: eukaryotic translation initiation factor 4B
FDR: false discovery rate
GSEA: gene set enrichment analysis
HILIC: hydrophilic interaction liquid chromatography
IGFBP5: insulin-like growth factor binding protein 5
MS: mass spectrometry
PCA: principal component analysis
polyP-22: polyP with chain length of 22 phosphate residues
polyP-45: polyP with chain length of 45 phosphate residues
polyP, PP: polyphosphate
RPS6KA1: ribosomal protein S6 kinase alpha-1
SRSF6: serine/arginine-rich splicing factor 6
TFA: trifluoroacetic acid
TiO_2_: titanium dioxide
TNC: tenascin C

