## Supplementary Figures and Legends for "Phosphoproteomic analysis of chondrocytes after short-term exposure to inorganic polyphosphate"

**Supplementary Figure 1 - Normalized value distributions**

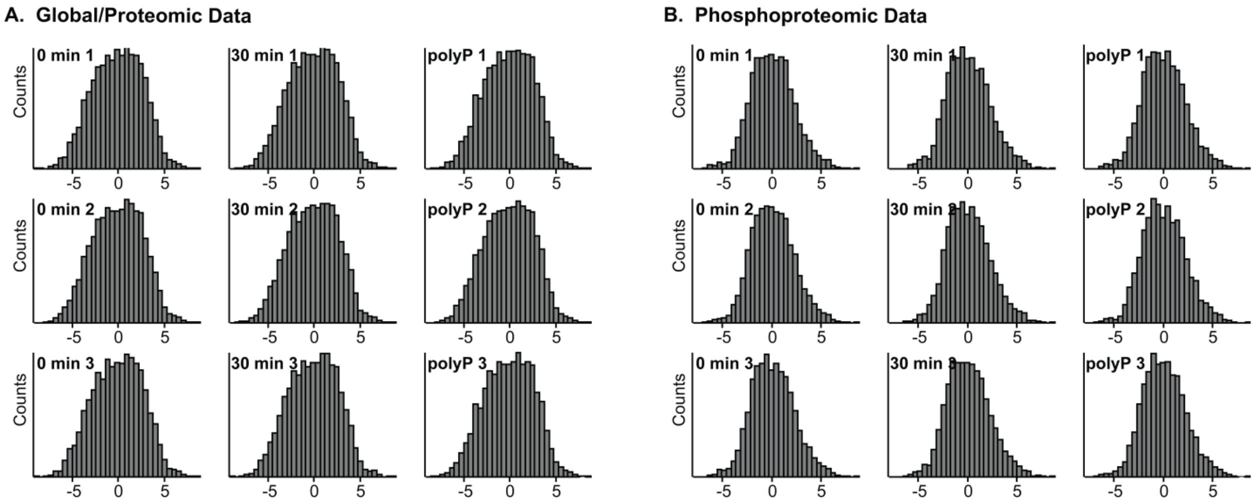


*Supplementary Figure S1. Distribution of normalized quantitative values of identified proteins (A) and phosphorylation sites (B) across different experimental conditions.*

Supplementary Figure 2 - Normalized value correlations

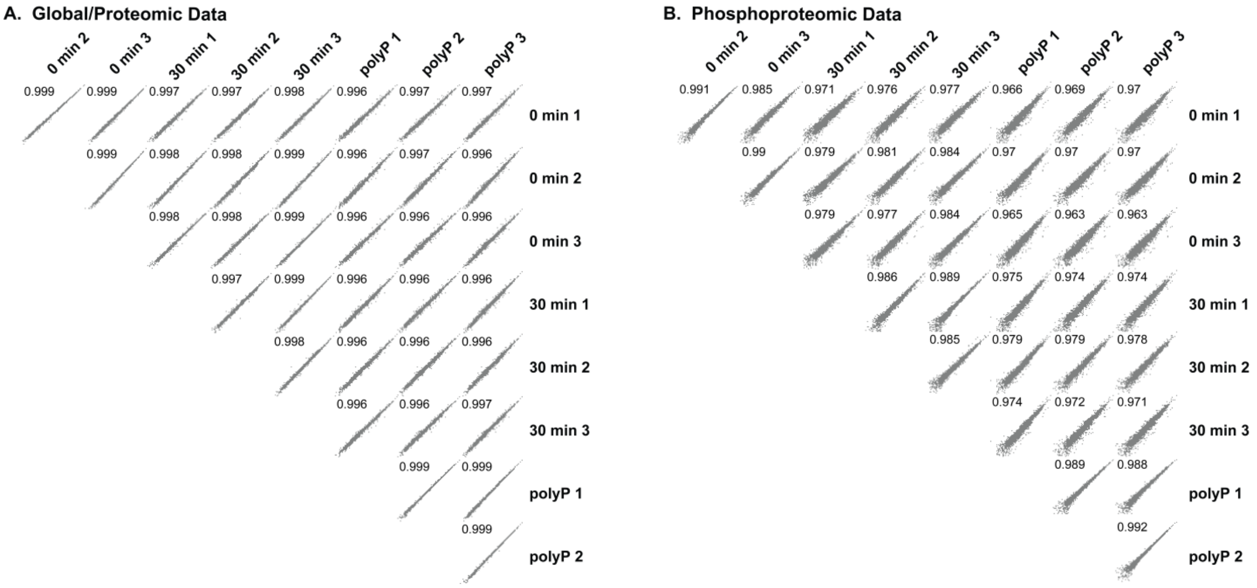


*Supplementary Figure S2. Correlation of normalized quantitative values of identified proteins (A) and phosphorylation sites (B) between individual samples in different experimental conditions with Pearson correlation coefficient inset in individual plots.*

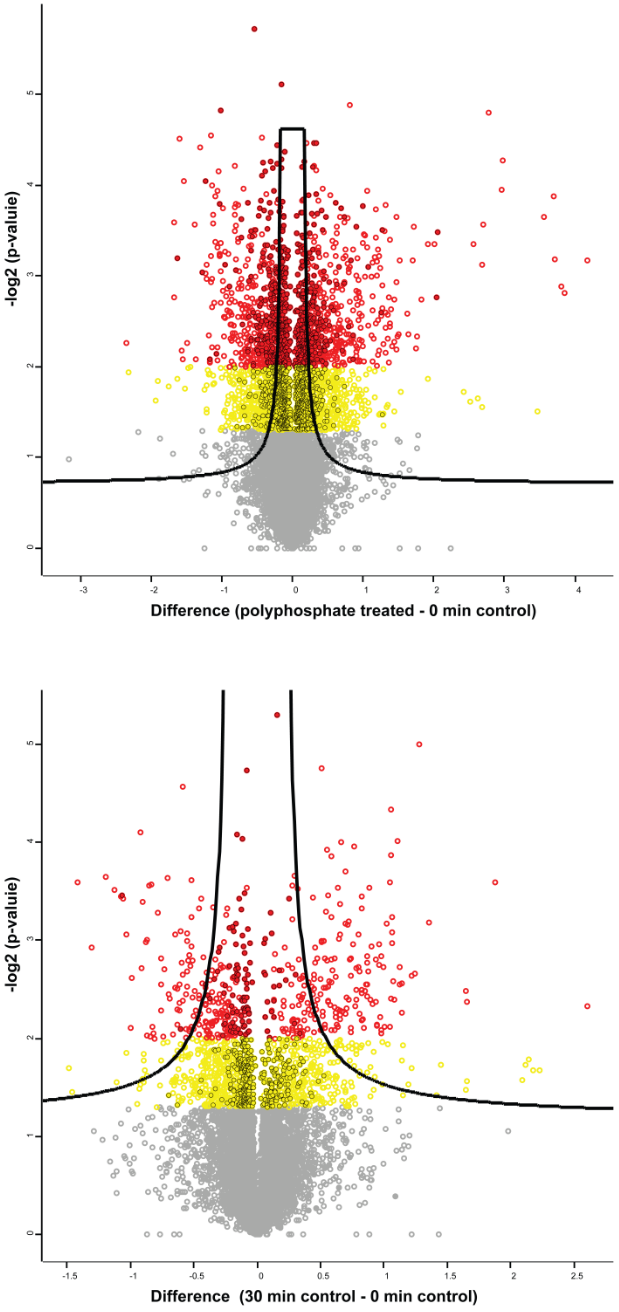


*Supplementary Figure S3.  Volcano plots representing the differences between polyP-treated and 0 min control (top) and 30 min post-media change and 0 min no media change control (bottom) merged data sets (protein and phosphorylation site quantitative information). IDs are represented by filled (proteins) and open (phosphorylation site) circles. The x-axis represents the difference in means. Student's t-test significantly altered proteins/phosphorylation sites are highlighted in yellow (p<0.05) and red (p<0.01). The highlighted lines represent an FDR cutoff (q<0.05) at an S0 = 0.1 artificial within group/sample type variance.*

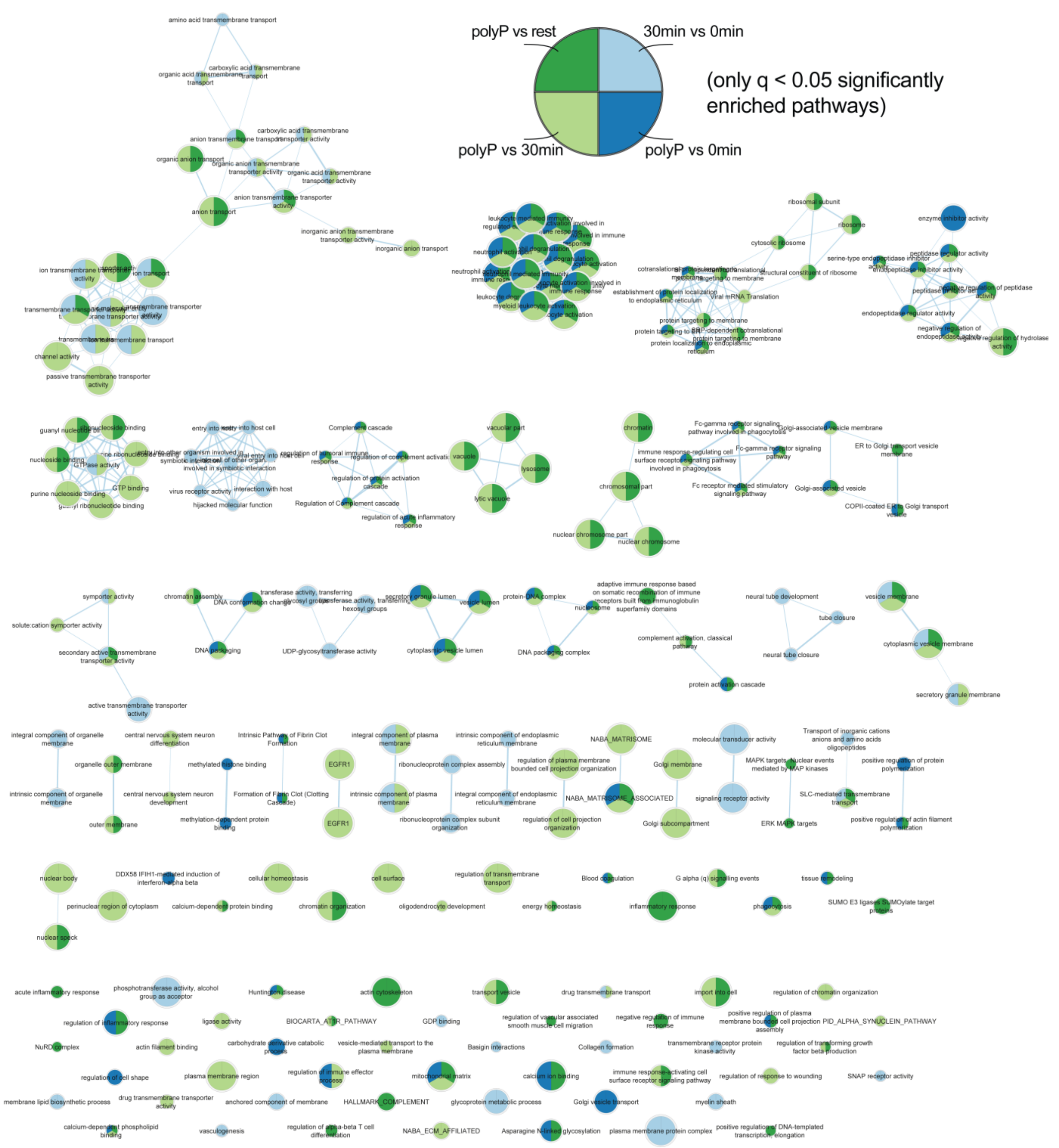


*Supplementary Figure S4A. Results of gene set enrichment analysis (visualized using the Enrichment Map plugin in Cytoscape). All significantly enriched gene sets identified in independent pairwise comparison analyses of different experimental conditions are visualized. Individual entries/nodes are coloured based significant enrichment (q<0.05) in each pairwise comparison. The thickness of the edges (lines) and the size of nodes (gene sets) represent the overlap between nodes found and the number of genes/proteins in the enriched gene sets in the data, respectively.*

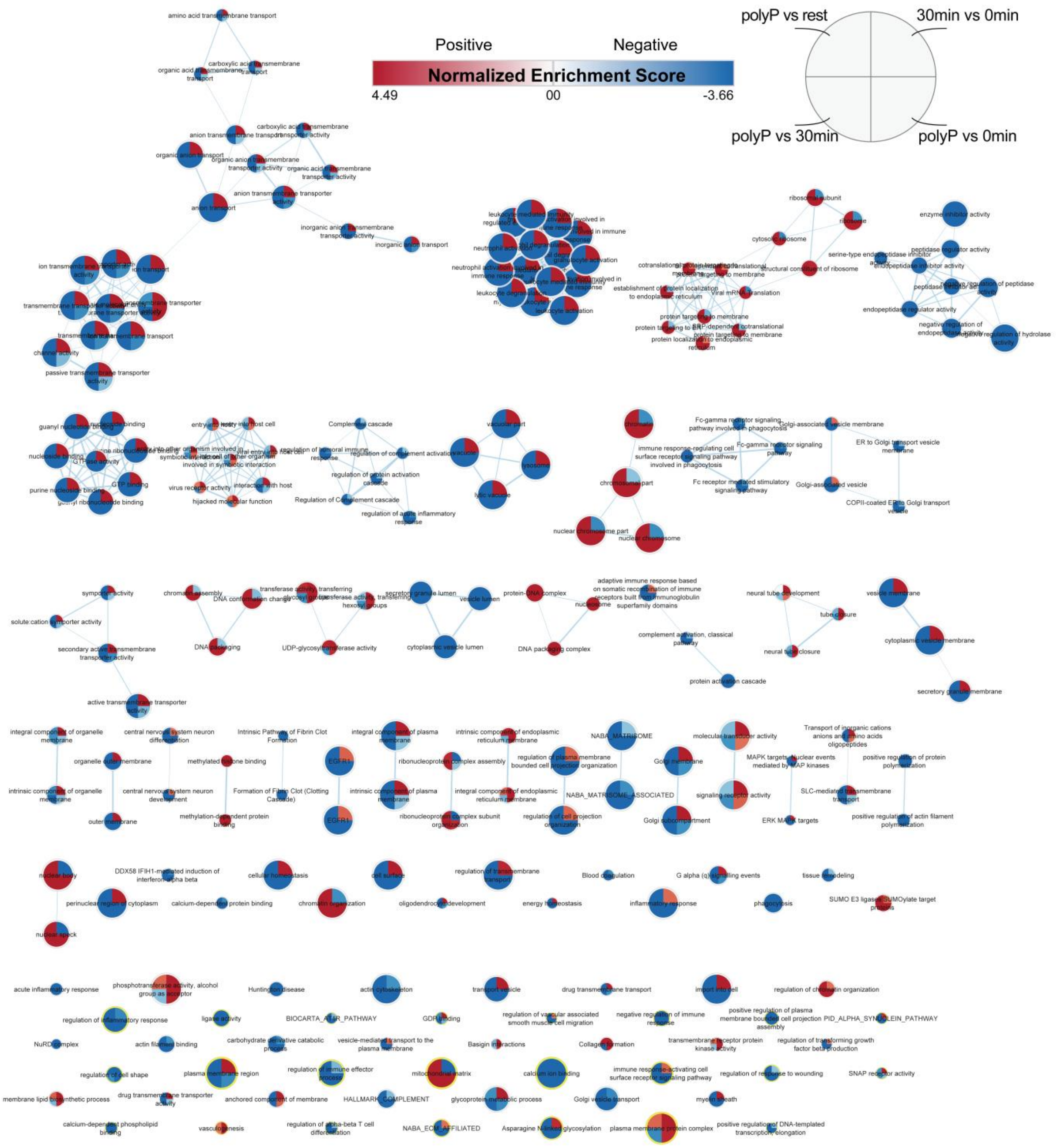


*Supplementary Figure S4B. Results of gene set enrichment analysis (visualized using the Enrichment Map plugin in Cytoscape). All significantly enriched gene sets identified in independent pairwise comparison analyses of different experimental conditions are visualized. Individual entries/nodes are coloured based on normalized enrichment scores (NES). The thickness of the edges (lines) and the size of nodes (gene sets) represent the overlap between nodes found and the number of genes/proteins in the enriched gene sets (nodes) in the data, respectively.*

*Supplementary Data S1. Combined results of proteomic and phosphoproteomic analyses. Raw and normalized quantitative data of polyphosphate treated, untreated, and media change control bovine chondrocytes at the protein and phosphorylation site level with associated column descriptions.*

*Supplementary Data S2. STRING analysis results. Full phosphoproteomic pathway analysis of 408 unique proteins based on significant changes in abundance in 1175 phosphosites in polyphosphate-treated versus non-treated bovine chondrocytes.*

*Supplementary Data S3. Data and Results for Statistical overrepresentation test in significant hits. Panther DB output of annotation coverage for all significantly altered (phospho)protein level identifications (from proteomic and phosphoproteomic analyses) for Gene Ontology (Biological Process, Cellular Component, Molecular Function) and Reactome Pathways. Separate analyses were performed for significantly up- and down-regulated proteins/phosphorylation sites.*

*Supplementary Data S4. all GSEA enriched gene set annotations in pairwise comparisons between the three experimental conditions (polyP-treated, untreated, and media change control) with associated enrichment score (ES), normalized enrichment score (NES), nominal P (NP), and false discovery rate (FDR) values.*
